# PSF broadening due to fluorescence emission

**DOI:** 10.1101/2019.12.21.885707

**Authors:** Jan Becker, Rainer Heintzmann

## Abstract

Fluorescent structures are nowadays commonly used in the field of biological imaging. However the emission of fluorescence is always given within a finite spectrum. When the imaging system’s point spread function is measured, this can be described as an incoherent summation of individual single-wavelength PSF’s, weighted by the emission spectrum. As the full-width-half-maximum of the PSF scales inversely with the wavelength and most emission spectra exhibit a fluorescence tail at longer wavelengths, this contributes significantly to lateral broadening. In our work we theoretically quantify this effect by deriving an analytic expression, which has been verified against some numerical simulations (rel. error on average 3 %). We report a FWHM-broadening on the order of 10 nm and additionally propose a way to overcome this broadening by splitting the emission spectrum into multiple wavelength segments. The corresponding image data is recombined in Fourier space by weighted averaging, leading to an improved signal-to-noise ratio at high spatial frequencies and a reduction of the FWHM of up to 8 nm (relative reduction of the broadening by ≈ 70 %). We also introduce a *corrected* wavelength, which in combination with already existing PSF calculation tools, describes a theoretical PSF which incorporates the aforementioned broadening effect.

## 1 Introduction

Many breakthroughs in the field of optical microscopy have been enabled due to the use of fluorescence [1, 2] as the utilized contrast mechanism. This is exemplified by many new imaging techniques which are commonly referred to as *super-resolution microscopy* [3-6], as all of them rely on fluorescence to break the diffraction limit [7].When quantifying the performance of an optical imaging system we usually use the concept of the point spread function (PSF), which acts as the impulse response of the corresponding system [8]. Its full-width-half-maximum (FWHM) is commonly regarded as a good measure for the imaging performance and scales inversely with the wavelength of light. However, every fluorescent structure emits light in a wavelength region, it’s emission spectrum. Most emission spectra show a tail towards longer wavelengths *(red* tail), which can be explained using the Franck-Condon principle [9] and Kasha’s rule [10]. When imaging such a fluorophore, the overall measurable PSF needs to be expressed as an incoherent sum of individual *single* wavelength PSFs, which all differ in FWHM, depending on the corresponding wavelength which is part of its emission spectrum (see Fig. 1b). We will show that this yields in a lateral broadening compared to the PSF estimated at the peak emission wavelength.

To our knowledge there is no quantitative description of this broadening effect yet. However, there is research on the influence of the illumination spectrum on spatial resolution of optical-coherence tomography (OCT) [11-14]. Their main goal is to inhibit the effects of PSF sidelobes in OCT by spectral reshaping of the used light source. The authors do not consider the effect of the emitted fluorescence but focus on changing the spectral properties of the illumination.

In the following we will first theoretically quantify the broadening effect and give an analytical expression. Next we will show the dependency of the broadening on the numerical aperture, the maximum emission wavelength and the shape of the emission spectrum. Thereafter this will be verified by numerical simulations. We are also proposing a new method to reduce the aforementioned broadening by splitting the emission spectrum in a finite number of wavelength segments and subsequent computational recombination of those, which will enhance the high spatial frequency signal-to-noise ratio (SNR). This is followed by a short discussion on our results and possible implications for future developments.

## 2 Theory

As described in sec. 1, the observable PSF *h(r)* at position *r* needs to be written as a incoherent sum:

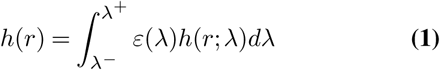

with ε(λ) the emission spectrum of the fluorophore, *h(r*;λ) the PSF for a single wavelength λ and λ^−^/λ^+^ the limits of the integration, given by the spectrum or a filter. We express ε(λ) as a *log-normal* distribution according to [15-17]:

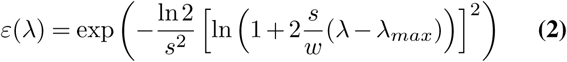

where λ_*max*_ is the wavelength corresponding to the emission maximum, *w* the FWHM of the spectrum and *s* a parameter indicating the shape or skewness of the emission spectrum. Note that for *s* → 0 the log-normal distribution reduces to a Gaussian (see sec. A; eq. 14). Typical examples of ε(λ) from three different fluorescent probes (Alexa-488, DAPI and mPlum; spectra from [18]) are shown in Fig. 1a. Equation 2 has been fitted to each dataset by minimizing the least-square error, yielding the corresponding parameters (Tab. 1). Fig. 1b shows the corresponding fit for DAPI in more detail.

**Table 1.**
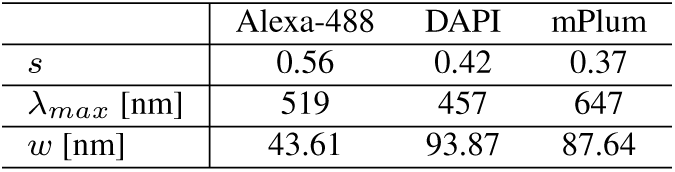
Results of fitting the log-normal distribution to the fluorescence spectra.

|  | Alexa-488 | DAPI | mPlum |
| --- | --- | --- | --- |
| $s$ | 0.56 | 0.42 | 0.37 |
| $\lambda_{max}$ [nm] | 519 | 457 | 647 |
| $w$ [nm] | 43.61 | 93.87 | 87.64 |

We introduce the constants:

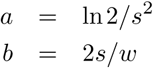

And rewrite equation 2 into:

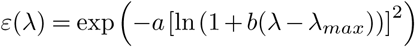

### A Description of the broadening effect

For the further analysis we approximate *h(r)* as a Gaussian. The best estimate (in least-squares sense) is given as [19]:

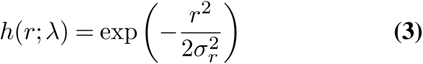

with:

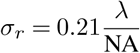

where NA refers to the numerical aperture of the detection objective. We express eq. 3 using a constant *C* = NA^2^/(2 ∙ 0.21^2^) and obtain:

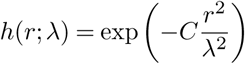

To analytically solve the integral given in equation 1, we write *h*(*r*;λ) as a Taylor series:

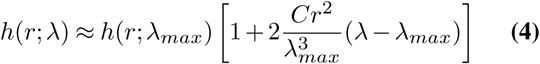

And introduce some constants *c*(*r*) and *d*(*r*):

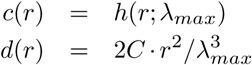

Hence equation 4 becomes:

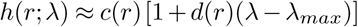

With this we are able to solve equation 1 and write:

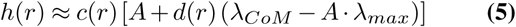

with the integrals:

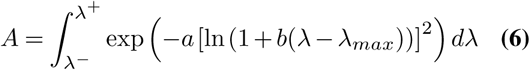

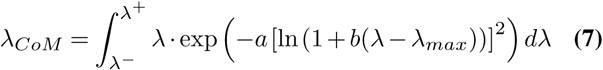

Note that *A* represents the area under the ε(λ)-curve, hence the total amount of emitted photons. λ_*CoM*_ is the wavelength which corresponds to the *center-of-mass* of the respective emission spectrum. Due to their positive skewness we find:

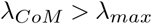

In appendix (sec. B) both integrals are solved, leading to:

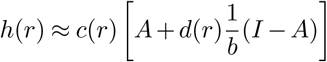

with an additional integral *I*, which result is given in eq. 15. For simplification we introduce another constant *D* = 1/*b* ∙ (*I/A* −1), yielding:

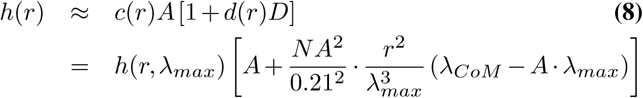

Note that equation 9 essentially tells us that the measured PSF *h(r)* can be expressed by the Gaussian approximation (given in eq. 3, with λ_*max*_), and a broadening term, which is given by a factor having the form of a Maxwell-distribution:

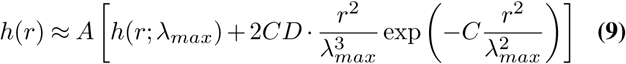

This is shown in Fig. 2. The black and green curves correspond to the Gaussian approximation at λ_max_ and the additional broadening represented by the Maxwell distribution. Adding these two function leads to our estimate *h*_*est*_.(*r*) indicated with the red (dashed) curve. The blue graph shows the simulated result accurately accounting for the PSFs at various wavelengths (more detail in sec. A), which is in good agreement with our theoretical findings. The inlet indicates the difference between our model and the detailed simulation. Even though our estimation fails at the sidelobes of the PSF, we are nevertheles still able to correctly estimate its FWHM.

### B Estimation of the broadening *B*

Next we want to approximate the FWHM of the broadend PSF, termed ∆*r*_*B*_. To do this, we first express eq. 9 using:

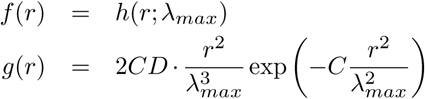

Therefore we write equation 9 as:

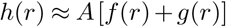

In the following we estimate the new FWHM (∆*r*_*B*_ = 2 ∙ *r*_1/2;*B*_) of the broadened PSF, as a linearly interpolated value between *r*_1/2_ and *r_max_.* A derivation and visualization of this procedure is given in sec. D. For the linear interpolation we need to parameterise a line given as: *y = m ∙ r +t.* The slope parameter *m* is given as (eq. 16):

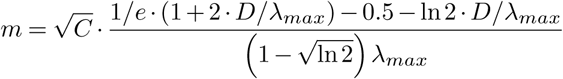

And the offset *t* as:

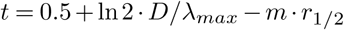

**Fig. 1.**
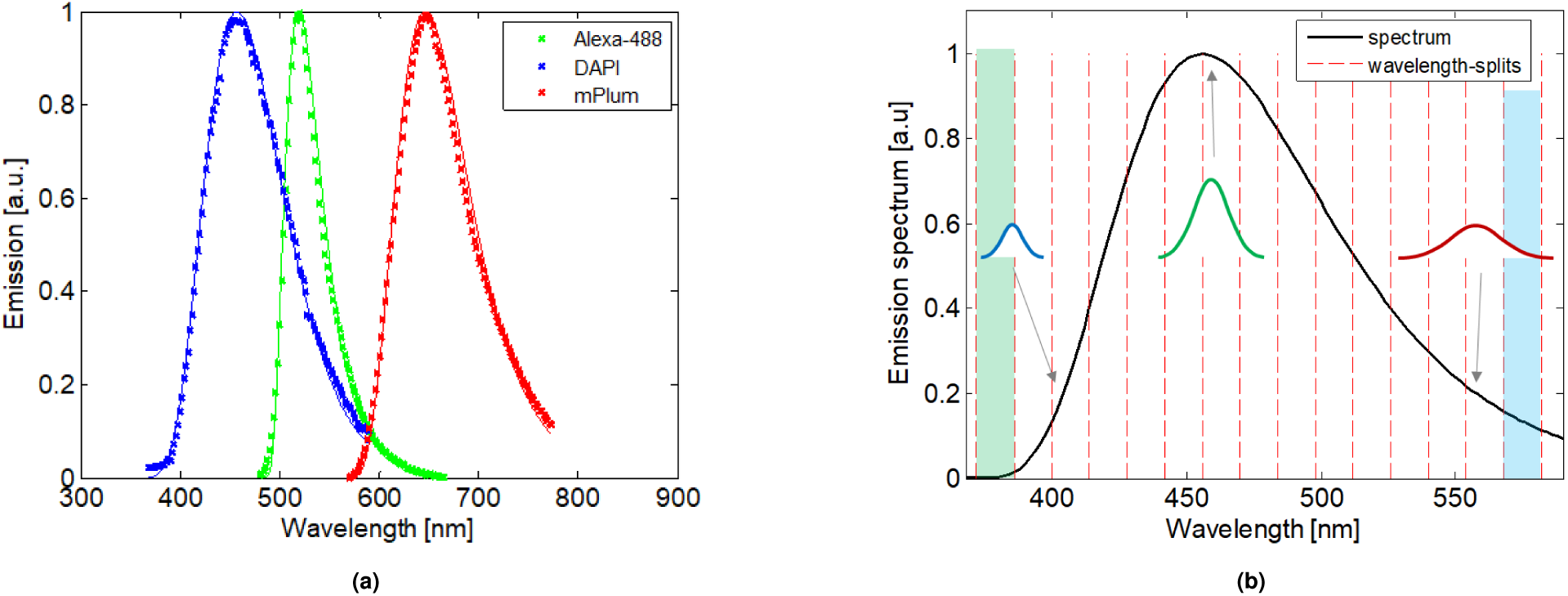
a) Fluorescence emission spectra of three different fluorophores (Alexa-488, DAPI and mPlum, data from [18]). A *log-normal* distribution has been fitted for each spectrum using a least-squares approach. b) Fitted spectrum of DAPI in more detail, including three Gaussians indicating the PSF broadening for longer wavelengths. The vertical lines represent the different wavelength bands in the splitting method, refer to sec. C.

**Fig. 2.**
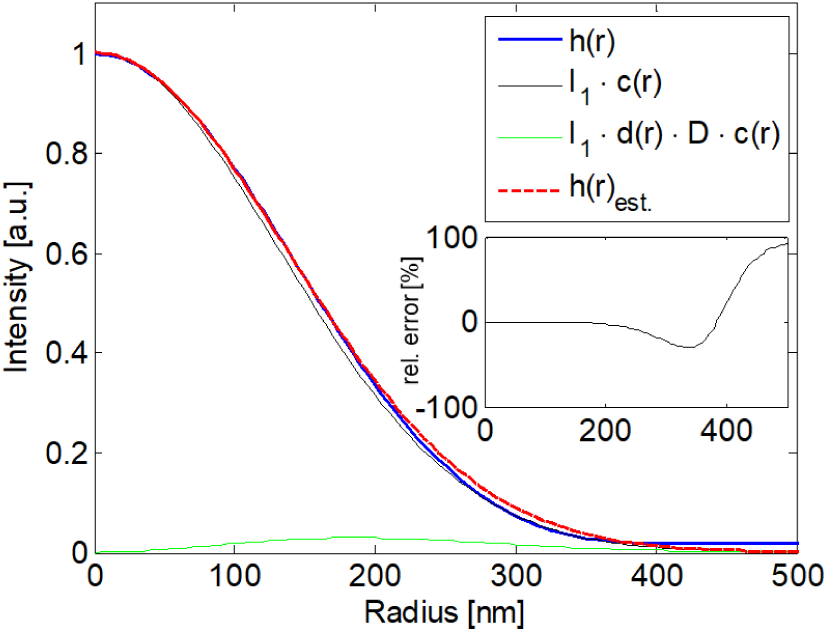
Broadened PSF as numerical result *h*(*r*) and theoretical prediction *h*_*est*_. (*r*). The inlet shows the relative difference [*h*(*r*) − *h*_*est*_(*r*)] /*h*(*r*) in percent. Strong deviations of our theoretical model and the numerical calculation only occur towards the sidelobes. The correct estimation of the FWHM is evident.

To obtain the new value *r_1/_*_2*;B*_ we need to solve *r = (y − t)/m* for *y* = 0.5. This leads to:

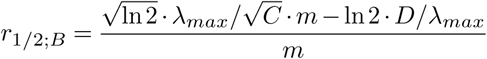

Yielding a new FWHM ∆*r*_*B*_ of:

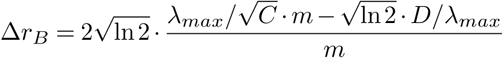

Comparing this to the previous FWHM value (∆*r* = 2 ∙ *r*_1/2_), yields an overall broadening *B* of:

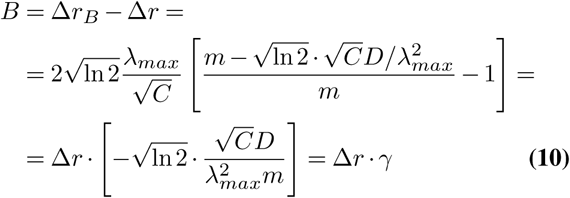

with:

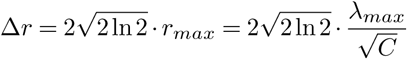

Finally we obtain a broadening by the relative factor γ:

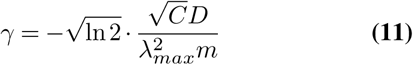

which is a function of:

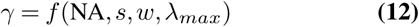

We are investigating the dependency of *B* on all the afore-mentioned parameters in section 3, where we will also compare our model to numerical simulations.

### C Wavelength splitting and recombination

A possible way to overcome the broadening as described in section A, is to split the emission spectrum into multiple wavelength bands. For each such band we capture an image and recombine these computationally, by weighted averaging in Fourier space (see appendix in [20]).

The different wavelength regions are marked in Fig. 1b with the vertical dashed red lines. In the depicted case the spectrum of DAPI has been segmented into 15 individual band. Each corresponding (sub-) image is described by the respective sub-PSF *h*_*i*_(*r*), which will differ in it’s width (depending on wavelength) and it’s maximum value (depending on the shape of the emission spectrum). After recombination we obtain an effective PSF, which can be related to an effective optical-transfer-function (OTF) [20], governed by the following equation:

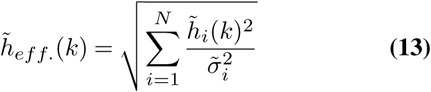

with 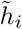 being the corresponding OTF of the *i*-th out of *N* total wavelength regions, *k* representing spatial frequencies and 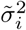 the variance associated with the *i*-th measurement.

## 3 Numerical results

In the following section we compare our model with numerical simulations and show the reduction of the broadening by wavelength splitting with subsequent computational recombination, leading to an improved SNR at high frequencies.

### A Comparing theoretical and numerical results

To verify the accuracy of equation 10, we compare our theoretical predictions to numerical results. To this aim we sim¬ulated PSF’s using the Richards & Wolf model [21] and ob¬tained the standard deviation σ, by least-squares fitting using a Gaussian. The corresponding FWHM is:

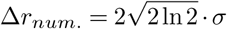

The calculations have been performed for two sets of optical parameters: NA = 0.8 / n = 1.00 (refractive index) and NA = 1.4 / n = 1.52. In the numerical case the broadening is obtained by replacing equation 1 by a discrete sum of the individual PSF’s *h*(*r*; λ) at the particular wavelengths within the spectrum. The respective PSF maxima are scaled by the corresponding value of ε(λ). The broadening follows as:

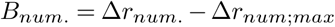

The FWHM of the broadend PSF is substracted from the FWHM of *h*(*r*;λ_*max*_). The difference of the two quantities is shown in Fig. 3. The inlet shows a zoomed region at the FWHM for both simulated and the theoretically estimated curve.

**Fig. 3.**
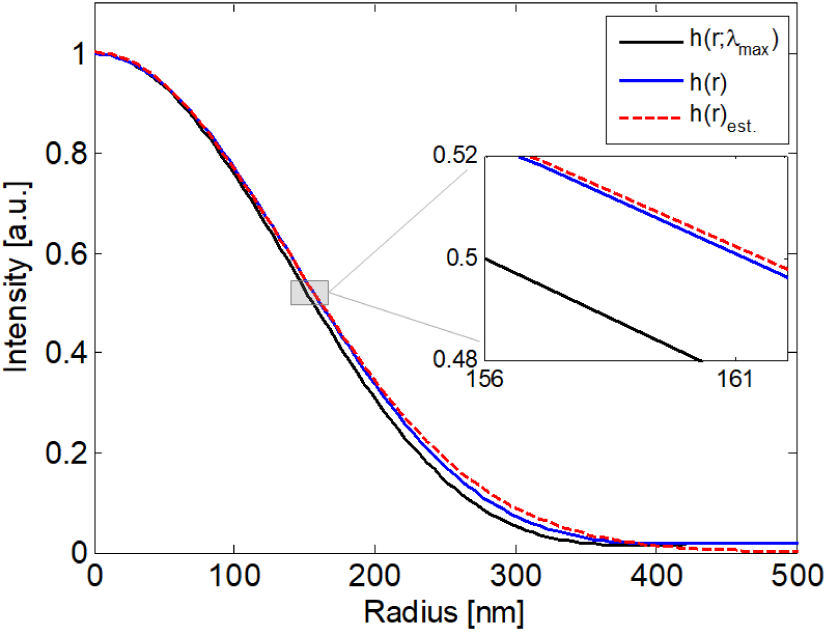
Comparison of the PSF with the peak emission only *h*(*r*;λ_*max*_) and the broadened PSF, estimated *h*_*est*_.(*r*) and numerically evaluated *h*(*r*). The inlet shows a zoomed region around the FWHM value and confirms our expectation of a lateral broadening.

Table 2 compares the numerical results *B*_*num*._ to the theoretical prediction *B*_*theor*._ (from eq. 10) for the different optical specifications.

**Table 2.** Comparing the broadening from the theoretical prediction (*B*_*theor*_.) with numerical results (*B*_*num*_.), for the three different fluorescent structures and two optical detection configurations (0.80 air & 1.4 oil objective). ∆*B* = *B*_*num*_. − *B*_*theor*_. in <u>nm</u>; relative difference in %.

| | NA / n | $B_{num.}$ | $B_{theor.}$ | $\Delta B$ | $\Delta B / B_{num.}$ |
| --- | --- | --- | --- | --- | --- |
| Alexa-488 | 0.80/1.00 | 10.79 | 11.04 | -0.25 | -2.32 |
|  | 1.40/1.52 | 6.41 | 6.31 | 0.10 | 1.60 |
| DAPI | 0.80/1.00 | 12.46 | 13.34 | -0.88 | 7.05 |
|  | 1.40/1.52 | 7.40 | 7.62 | -0.22 | -3.02 |
| mPlum | 0.80/1.00 | 11.22 | 11.19 | 0.03 | 0.27 |
|  | 1.40/1.52 | 6.62 | 6.39 | 0.22 | 3.38 |

Both, numerical and theoretical, results show a broadening on the order of 10 nm. The observed deviation is well below 1 nm accuracy (rel. error on average is 3 %), showing that our theoretical analysis gives good predictions. Tab. 2 also indicates the inverse proportionality between broadening and NA.

### B Dependency of *B* on various parameters

As noted already in equation 11, the broadening depends on four parameters: the numerical aperture NA, the shape parameter s, the FWHM *w* of the emission spectrum and the wavelength of maximum emission λ_*max*_. We have investi-gated the dependency on each of these parameters by solving equation 10 numerically, the results are shown in Fig. 4a.

**Fig. 4.**
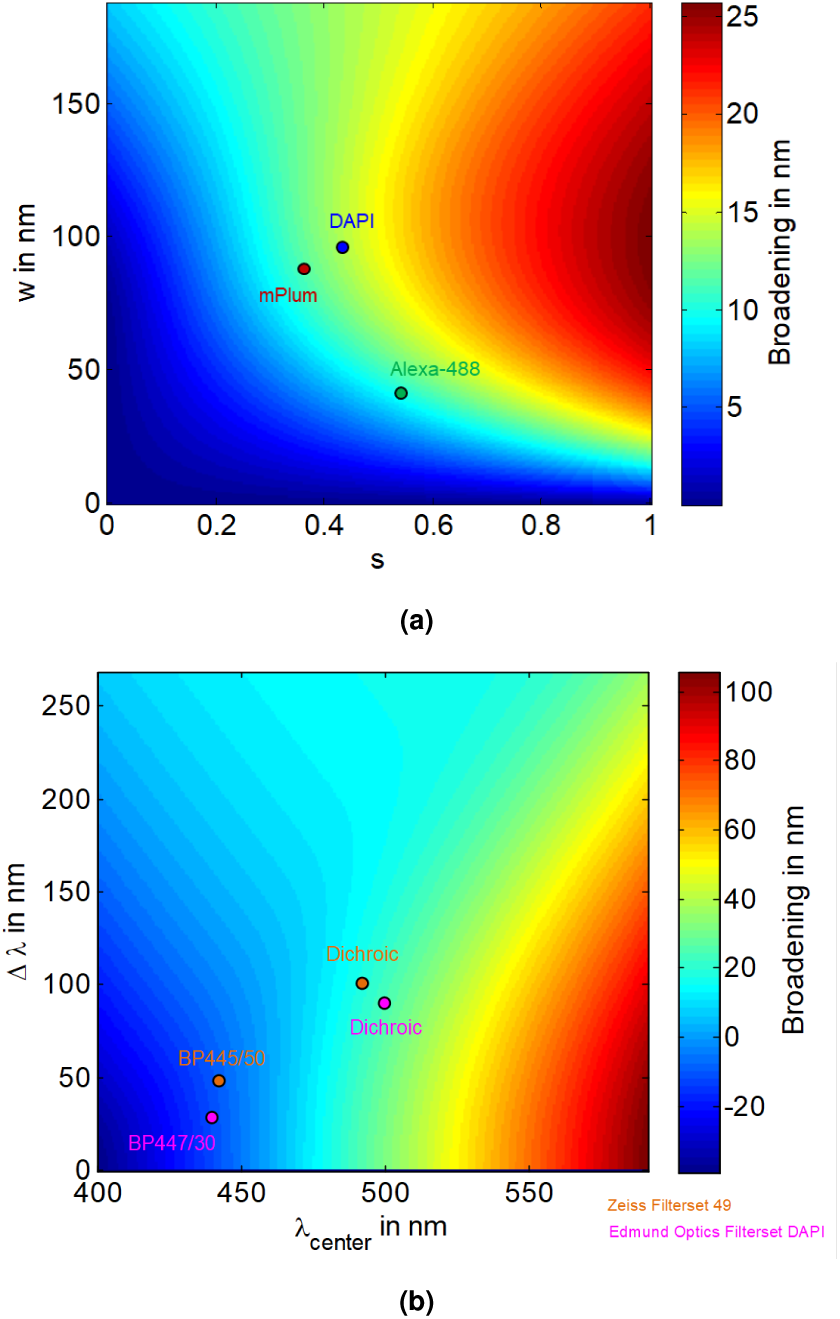
The broadening (given in nm), depending on the shape of the emission spectrum (a) and the bandpass characteristics (b) of an additionally used emission filter. Wider emission spectra and more asymmetry leads to a stronger PSF broadening. Choosing a narrow bandpass filter reduces the broadening, but comes at the expense of the total number of detected photons.

Fig. 4a shows the dependency of the broadening on the shape of the emission spectrum. It can be seen that mainly the asymmetry, indicated by *s >* 0, contributes to the aforementioned broadening effect. Wider emission spectra (larger *w*) also lead to a stronger broadenig, however the influence is not as prominent as the shape parameter.

Most imaging applications do not make use of the complete emission spectrum. Usually a fluorescence filterset consists of an excitation, a dichromatic beamsplitter and an emission filter. Latter is typically a bandpass (BP) with a certain center wavelength λ_*center*_ and a bandwith ∆λ. Fig. 4b shows the dependency of the broadening on those two parameters, when simulating DAPI according to the values found in Tab. 1. The stronger influence is noticeable for increasing the center wavelength, compared to only using a wider BP. For narrow filters we even observe a negative broadening: *h*(*r*) *< h*(*r*;λ_*max*_). However, decreasing the spectral width comes at the cost of a reduced total number of detected photons, hence overall SNR.

### C Simulation of wavelength splitting & recombination

Our proposed method to overcome this effect is shown here for a splitting of the spectrum of DAPI into *N* =15 segments, as shown in Fig. 1b. We investigate the imaging using a NA = 0.8 air-objective. The individual PSF’s are recombined, yielding an effective PSF, which is given as the inverse Fourier transform of equation 13. We show the resulting OTF in Fig. 5a in *log*-scale. The cyan and green colored curves correspond to the sub-OTF’s of the first and 15^*th*^ segment.

**Fig. 5.**
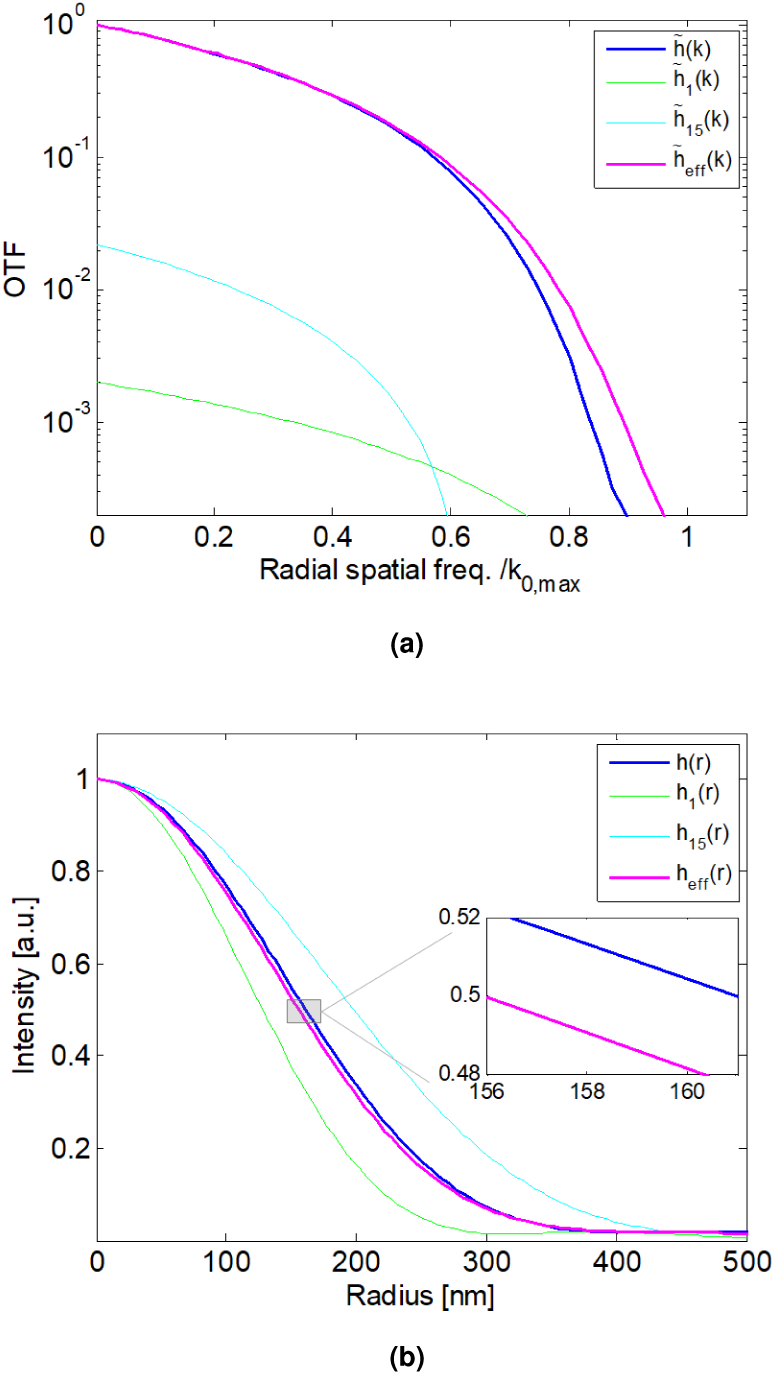
a) OTF for the different wavelength bands 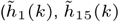; marked in Fig. 1b, the broadened 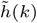 and the recombined 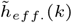 result in *log-scale*. The different OTF’s are efficiency-normalized, hence their value for *k* = 0 scales with the number of photons available in that wavelength region. Note that their respective cut-off frequency differ (*k*_0_,_*max*_ = cutoff of lowest emission wavelength). b) PSF of the different wavelength regions from Fig. 1b (*h*_1_(*r*), *h*_15_(*r*)) and the recombined result *h*_*eff*_ (*r*). Note the reduced FWHM compared to the (broadened) PSF without splitting *h*(*r*).

Fig. 5a indicates that the cut-off frequency depends on the wavelength region used for imaging. Larger wavelengths (cyan graph) lead to a reduction in transferable spatial frequency information. After splitting and recombination a clear improvement, compared to the (broadend) result, is noticeable. Especially closer to the high spatial frequency, which is due to the fact, that the weighted averaging recombination approach optimizes the SNR [20] which becomes crucial towards higher frequencies. The corresponding PSF-graphs are shown in Fig. 5b, indicating the reduction in FWHM. Numerical values for both, broadend and recombined, PSF’s are given in Table 3.

**Table 3.** Comparing the FWHM of the broadend PSF (∆*r*_*B*_) with the (effective) recombined result of the wavelength splitted data (∆*r*_*eff*_.), for the three different fluorescent structures and the two optical detection configurations. ∆_*r*_ = ∆*r*_*B*_ − ∆*r*_*eff*._ in <u>nm</u>; relative broadening in %.

| | NA / n | $\Delta r_B$ | $\Delta r_{eff}$ | $\Delta_r$ | $\Delta_r / B_{num.}$ |
| --- | --- | --- | --- | --- | --- |
| Alexa-488 | 0.80/1.00 | 356.94 | 353.40 | 3.54 | 32.81 |
|  | 1.40/1.52 | 211.92 | 210.39 | 1.53 | 23.87 |
| DAPI | 0.80/1.00 | 317.26 | 309.91 | 7.35 | 59.00 |
|  | 1.40/1.52 | 188.35 | 184.26 | 4.08 | 53.54 |
| mPlum | 0.80/1.00 | 443.73 | 435.66 | 8.07 | 71.93 |
|  | 1.40/1.52 | 263.61 | 260.59 | 3.02 | 45.62 |

The reduction in FWHM on average is 4.6 nm, but also depends strongly on the shape of the emission spectrum. The maximum improvement can be observed for *mPlum*, which spectra shows the lowest skewness parameter s. Therefore, depending on the fluorescent structure used for imaging, our wavelength splitting approach might yield considerable enhancements of up to 8 nm (relative change of 72 %).

**Fig. 6.**
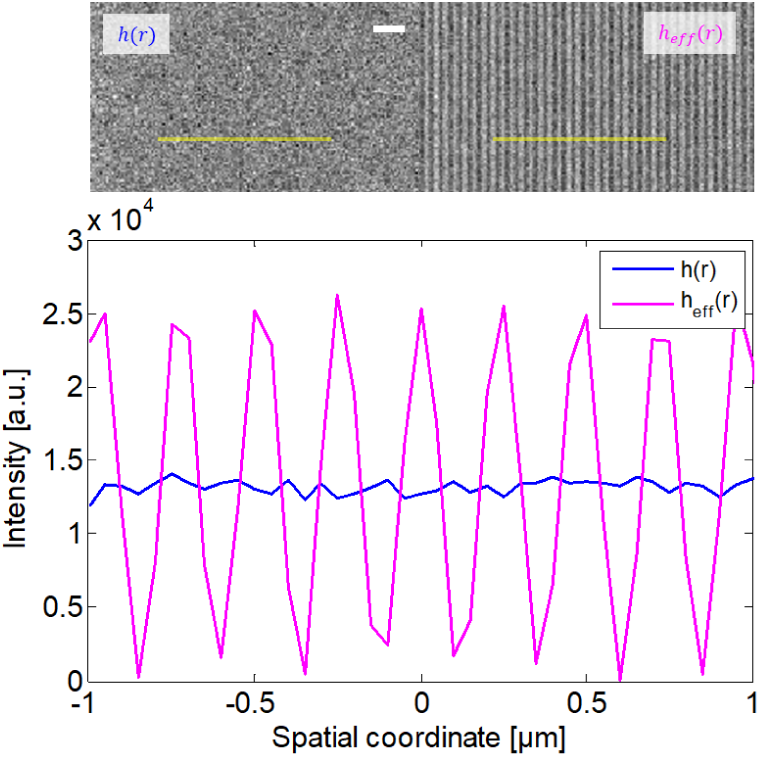
Top: Simulation results of imaging with (magenta) and without (blue) wavelength-depending splitting (*percentile-scaling*). The high frequency line pattern can only by observed when the splitting technique is used (due to the improved SNR). Scale bar = 2 γm. Bottom: line profile indicating the gain in modulation depth.

To illustrate the possible enhanczement of our new imaging approach, we show simulations of a fine grid objects in figure 6. Which has been convolved with the respective PSF (*h*(*r*) or *h*_*eff*_ (*r*)). Poisson noise has been applied to the simulated data, with a 1000 photons in the maximum pixel.

For Fig. 6 the frequency of the line pattern has been raised until the broadend image result only shows noise (top). Whereas the recombined result is still able to clearly resolve the given line pattern (bottom). This comes from the aforementioned strong increase in high frequency SNR.

## 4 Discussion

In the derivation of our theoretical expression for the broadening *B*, we assumed that the emission spectrum can be expressed as a log-normal distribution. Of course there exist a large variety of spectral shapes in real life [18]. Our theory is only applicable to those, who resemble the suggested log-normal curve. However, the shown results are still useful to give a first impression on the order of magnitude the *red tail fluorescence PSF broadening effect* might have in their specific imaging task. Our Gaussian expression of the PSF, despite being an approximation yielded relatively precise results for the FWHM as shown in Fig. 2. Only towards the sidelobes, which is far beyond the FWHM point, our approximation clearly fails. In terms of searching for the lowest PSF broadening, our theoretical results show that a symmetric (*s* → 0) and narrow (*w* → 0) emission spectrum (Fig. 4a is crucial. This seems expectable as it refers, in the limiting case, to only a single wavelength being emitted. Which of course will lead to no broadening at all as eq. 1 only consists of a single contribution. We have added markers for the three examined fluorphores into Fig. 4a. It becomes evident that there might exist fluorescent markers which show a much stronger broadening, than those we chose. Our analytical description helps in identifying those. As has been shown in section B, the overall broadening also depends on the emission filter used. We are comparing to filtersets, common for imaging DAPI (Zeiss Filterset 49 + Edmund Optics Filterset DAPI). The scenario of only using an excitation + dichroic filter has been marked in Fig. 4b with the labels "Dichroic". The broadening, for both filtersets is in the region of 10 − 15 nm. Adding the BP-filters (BP445/50 + BP447/30) leads to a drastic reduce of the broadening (*B* actually becomes < 0). As already mentioned earlier, this comes at the cost of a reduced number of detectable photons, as only a small part of the emission spectrum is used for imaging. The broadening decreases towards higher numerical aperture (Fig. 4a), but so does the overall FWHM. In fact, the relative broadening (= *B/*∆*r)* is constant at 4.76 % (see Fig. 8a. Going for longer maximum emission wavelengths (λ_*max*_) slightly decreases the broadening (Fig. 8b). We also found that already existing PSF calculation tools, which usually take λ_*max*_ as their input, can be modified to also include the aforementioned broadening effect by taking the substitution λ_*max*_ → λ_*corr.*_ with:

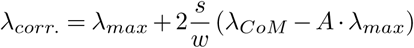

Yielding in a wavelength which lies between the maximum emission and the "center of mass" of the spectrum. The following table compares the broadend FWHM ∆*r* with the corrected values ∆*r_corr_.*.

We find that using the corrected wavelength reduces the broadening to on average 1.6 nm, which corresponds to an improvement of one order of magnitude. However, we do notice a strong dependency on the emission shape of the specific fluorophore, as DAPI almost shows no broadening at all. Including this information in already exisiting PSF-models might give some benefits in all methods were an accurate description ofthe PSF is crucial, e.g. image reconstruction based instruments like ISM [22, 23]. Our wavelength splitting approach shows a clear enhancement of SNR at high spatial frequencies. We assumed to work in the photon-limited regime, hence neglect additional noise components coming from the detectors. As advancement in camera technology has brought the read-out noise down to a negligible value of ≤ 1 electron RMS [24]. As noted before, in our simulation we have split the emission spectrum of DAPI into 15 segments. This seems to be more challenging in a real experimental setup. The most straightforward way to implement the wavelength splitting approach, would be to use several bandpass filters which are positioned successively. However, there already exist spectral microscopes capable of imaging a given field-of-view, while also recording the corresponding spectra pixel by pixel [25]. Such devices, e.g. a spectral confocal microscope, could be easily used to generate the data needed for our wavelength splitting approach. As the enhancement mainly can be seen at high spatial frequencies, it is clear that the most suited application of this technique is in the detection at or near the resolution limit. A similar enhancement in effective resolution can also be achieved by using a detection objective with higher NA. However when the highest available NA is not enough to resolve some high frequency structures, e.g. for refractive index matching when imaging biological sample, our method can still precede.

**Table 4.** Comparing the FWHM of the broadend PSF (∆*r*_*B*_) with the theoreti-cal calculation (∆*r*_*corr.*_) using the corrected wavelength λ_*corr.*_. The corrected broadening value is: *B*_*corr.*_ + ∆*r*_*B*_ − ∆*r*_*corr.*_ in <u>nm</u>; rel. broad. in %.

| | NA / n | $\Delta r_B$ | $\Delta r_{corr.}$ | $B_{corr.}$ | $B_{corr.}/B_{num.}$ |
| --- | --- | --- | --- | --- | --- |
| Alexa-488 | 0.80/1.00 | 356.94 | 360.88 | -3.94 | -36.51 |
|  | 1.40/1.52 | 211.92 | 214.22 | -2.30 | -35.92 |
| DAPI | 0.80/1.00 | 317.26 | 316.86 | 0.40 | 3.24 |
|  | 1.40/1.52 | 188.35 | 188.10 | 0.25 | 3.38 |
| mPlum | 0.80/1.00 | 443.73 | 441.89 | 1.84 | 16.39 |
|  | 1.40/1.52 | 263.61 | 262.52 | 1.09 | 16.48 |

## 5 Conclusion

We have found an analytical expression describing the effect of PSF broadening due to the finite emission spectrum of the fluorescent markers used in imaging. Our investigation shows that the broadening for typical fluorphores is on the order of 10 nm, hence not negligible in every application. It is important to understand that his broadening effect is fundamental to fluorescent imaging. It is expected that an experimentally obtained PSF differs from the theoretical counterpart. Often this is attributed to aberrations or misalignment of optical components. But even with the perfect system, in the noise-free case, some broadening will be observable due to the effect described in our work. Additionally we have introduced a technique which is able to overcome this broadening, without compromising the SNR. The proposed wavelength splitting approach uses the fact that computational recombination is able to use the image information more effectively.

## Acknowledgements

The style format of this publication is based on https://www.overleaf.com/latex/templates/henriqueslab-biorxiv-template/nyprsybwffws, which has been manipulated according to our needs.

## Authors contributions

J.B. conceived the initial idea, worked on the theoretical details and performed the numerical investigation. R.H. took care of funding, provided the research environment and supervision. Both (J.B. and R.H.) worked on the manuscript.

## Appendix

### A Log-normal distribution for *s* → 0

Our model of the emission spectrum is given as:

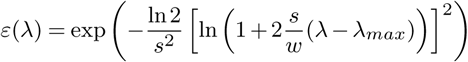

The exponent can be rewritten into:

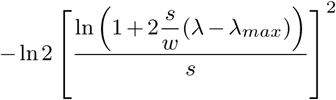

Calculating the limit of the term in brackets for *s* → 0 gives:

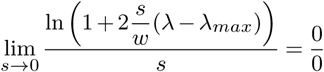

We need to apply the rule of l’Hopital yielding:

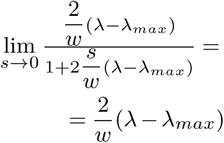

Taking this, we obtain the exponent again by:

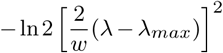

Relating the FWHM *w* to the standard deviation *σ*_*ε*_ of the Gaussian distribution:

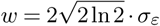

Finally yielding in:

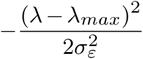

Showing that in the limit of vanishing "skewing" parameter s, our expression for the fluorescent emission becomes:

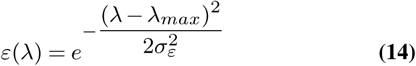

### B Solving integrals *A* and λ_*CoM*_

Equation 7 can be solve by substituting *u* = 1 + *b*(λ − λ_*max*_) (*du/d*λ = *b*; λ = λ_*max*_ + (*u* − 1)/*b*), shown here for *I*_1_:

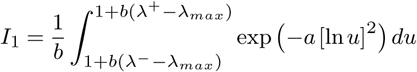

An additional substitution of *v* = ln[*u*] (*dv*/*du* = 1/*u*; *u* = *e*^*v*^) yields:

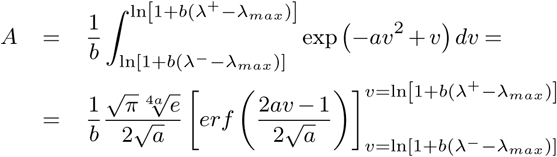

with *er f*(...) being the (Gauss) error-function. Note that *A* neither depends on *r* nor on λ.

The same (substitution) procedure can be done for λ_*CoM*_:

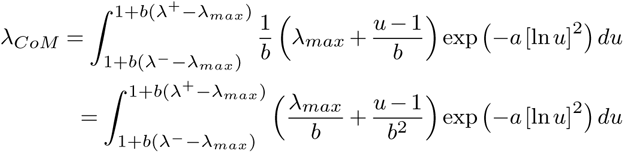

Leading to:

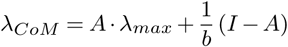

with an additional integral *I* to solve:

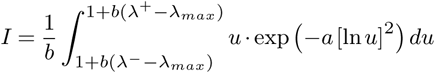

### C Solving integral *I*

Integral *I* is solved by the same substitutions which have already been used in section B.

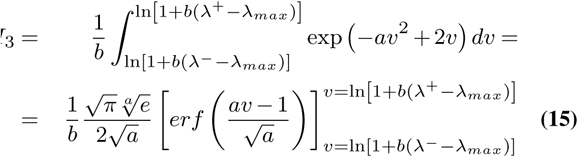

### D Values of *f*(*r*) and *g*(*r*) at *r*_1/2_ and *r*_*max*_

For estimating the broadend FWHM ∆*r*_*B*_ we are employing a linear interpolation as shown in Fig. 7.

**Fig. 7.**
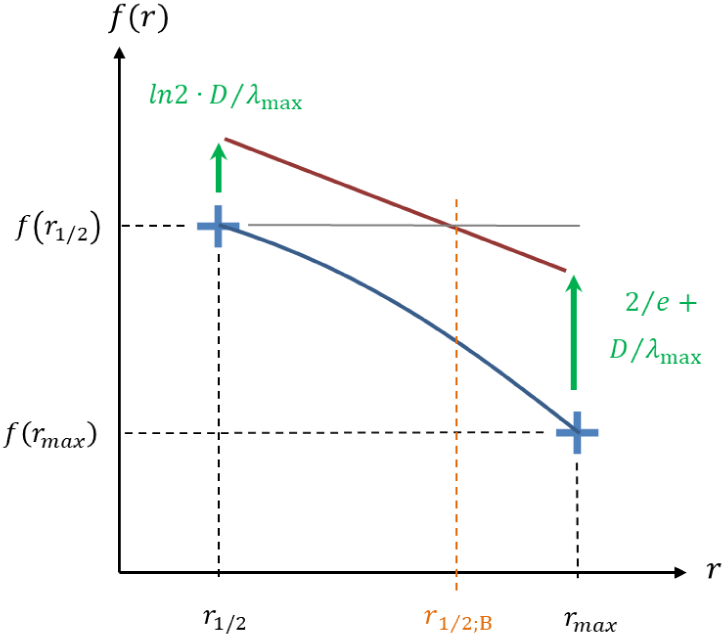
Scheme to estimate the broadened FWHM 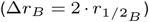 as a linear interpolation (red line). Green shows the additional contributions which will lead to the broadening.

As indicated, corresponds to *f* (*r*_1/2_) = 0.5 and *r*_*max*_ to the position of the maximum of *g*(*r*). For the interpolation we use the two points (*r*_1/2_, *f*_1/2_ + *g*_1/2_) and (*r*_*max*_,*f*_*max*_ + *g*_*max*_). Hence we can define the slope parameter *m* as:

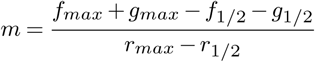

First we compute the position of the maximum *r*_*max*_ of *g*(*r*). To do this we obtain the first derivative as:

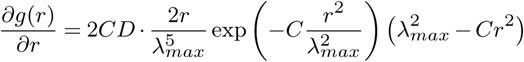

Setting this equal to zero and solving for *r* yields:

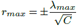

With this we are able to obtain the maximum value of *g*(*r*) as:

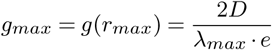

The corresponding value for *f*(*r*) is

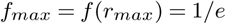

Now we are looking for the point at which *f*(*r*) has half of it’s maximum value:

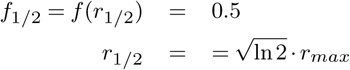

The corresponding value for *g*(*r*) at this position is:

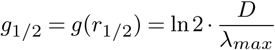

Which finally leads to:

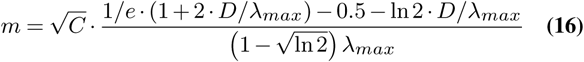

### E Broadening *B* as a function of *NA* and λ_*max*_

The dependency of the broadening on the NA and λ_*max*_ can be seen in Fig. 8a and Fig. 8b respectively.

**Fig. 8.**
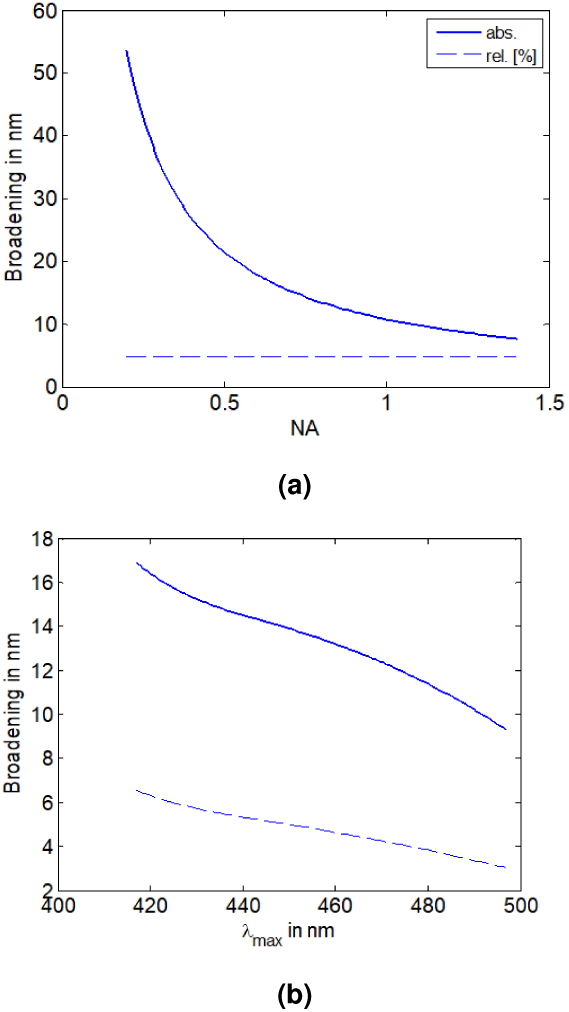
Broadening (in nm) in dependency of the detection numerical aperture a) and the emission maximum wavelength λ_*max*_. As already described in sec. B, the relative broadening (*B*/∆*r*) shows a linear dependency (a: constant at 12.84 %, b: declining line). Overall the relative broadening is on the order of 10%.

When changing the numerical aperture from 0.2 to 1.4, a reduction of the broadening by a factor of ≈ 86 % can be observed. The relative broadening (*B*/ λ*r*) however stays constant at 4.76 %.

In case of increasing the maximum emission wavelength of the fluorescent marker, also a decrease in broadening is observed. Interestingly the behaviour is not strinctly linear.

